# A Patient-Derived Scaffold-Based 3D Culture Platform for Head and Neck Cancer: Preserving Tumor Heterogeneity for Personalized Drug Testing

**DOI:** 10.1101/2025.07.31.667921

**Authors:** Alinda Anameric, Emilia Reszczyńska, Tomasz Stankiewicz, Adrian Andrzejczak, Andrzej Stepulak, Matthias Nees

## Abstract

Standard 3-D models for head-and-neck cancer (HNC) often lose stromal partners that influence drug response or never include them. We developed a patient-derived cell culture system that maintains tumor cells, cancer-associated fibroblasts (CAFs), and cells undergoing partial epithelial-to-mesenchymal transition (pEMT) for rapid sensitivity testing. Biopsies from four HNC patients were enzymatically dissociated. CAFs were directly cultured, and their conditioned medium (CAF-CM) was collected. Cryopreserved tumour cell suspensions were later revived, screened in five different growth media in 2-D conditions, and the most heterogeneous cultures were re-embedded in 3-D hydrogels with varied gel mix, medium, and seeding geometry. A perimeter-derived complexity index was used to quantify tumoroid morphology and viability after exposure to cisplatin or the Notch modulators RIN-1 (activator) and FLI-06 (inhibitor), which were assessed by live imaging and WST-8 assays. ECM-2 medium alone produced compact CAF-free spheroids, whereas ECM-2 supplemented with CAF medium generated invasive aggregates that deposited endogenous matrix; Matrigel plus this medium and single-point seeding yielded the highest complexity scores. Two of the three patient tumoroids were cisplatin-sensitive, and all showed significant growth inhibition with the FLI-06 inhibitor, while RIN-1 induced minimal change. The optimised scaffold retains tumour–stroma cross-talk and affords patient-specific drug-response data within days, supporting personalised treatment selection in HNC.

## Introduction

Head and neck cancer (HNC) is the seventh most common cancer globally, resulting in 325,000 deaths per year. Tobacco, alcohol abuse, and oncogenic viruses, including human papillomavirus (HPV), are the main risk factors in HNC [1]. With an incidence of over 90%, the most common tumor type is head and neck squamous cell carcinoma (HNSCC), which originates from the mucosal epithelium in the oral cavity, pharynx, larynx, nasal cavity, and salivary glands. In contrast, adenoid cystic carcinoma (ACC) comprises only 1–2% of cases. ACC is typically a salivary gland malignancy, and its occurrence at the base of the tongue is considered rare [2]. The complex relationship between the squamous cancer cells, the tumor microenvironment (TME), and the extracellular matrix (ECM) provides a broad spectrum for potential therapeutic interventions, regardless of the clinical stage [3].

Cancer-associated fibroblasts (CAFs) are an essential component of the TME. The diversity of CAFs contributes significantly to the intra- and inter-tumor heterogeneity of HNSCC. Recent research showed that CAFs comprise at least 3 or 4 functionally different subtypes: myofibroblastic CAFs (myCAFs), inflammatory CAFs (iCAFs), vascular CAFs (vCAFs), and antigen-presenting CAFs (apCAFs) [4]. The most frequent myCAFs can be further subdivided into C1-type myCAFs, characterized by low expression of alpha smooth muscle actin (IF) and high bone morphogenetic protein 4 (BMP4), and C2-type myCAFs, which show the opposite pattern [5]. The intra- and inter-tumor heterogeneity of HNSCC also includes different epithelial phenotypes. One of these displays both epithelial and mesenchymal traits, undergoing a phenotypic shift described as partial epithelial-to-mesenchymal transition (pEMT). Recently, four major epithelial clusters were identified in HNSCC: (1) Sox2-high/K14-low cells with high density, (2) K14-positive large cells lacking pEMT and stem cell markers, (3) a stem-cell-like cluster expressing K14, Sox2, and Bmi1, and (4) cells co-expressing K14, Slug, and Vimentin [6].

Cisplatin is a standard chemotherapeutic drug used in HNSCC treatment, while advanced or metastatic cases (RM-HNSCC) often receive combination therapy with docetaxel, cisplatin, and 5-fluorouracil. Targeted treatments include monoclonal antibodies like cetuximab and bevacizumab, and pathway-specific inhibitors such as temsirolimus and rapamycin targeting mTOR signaling [7]. There are also Notch signaling modulators under pre-clinical investigation, such as RIN-1, which activates Notch via RBPJ inhibition [8], whereas FLI-06 acts as a potent Notch inhibitor [9].

3D cell culture systems overcome the limitations of two-dimensional models by better replicating the morphological and functional features of the tumor microenvironment. 3D cultures can be based either on scaffold-free or scaffold-based systems. Scaffold-free systems promote the spontaneous formation of multicellular aggregates or organoids and are typically promoted by non-adherent and suspension-culture techniques. In contrast, scaffold-based cultures typically rely on synthetic (ceramics, metals, or polymers) or natural scaffolds (polysaccharides, fibrous proteins, ECM-derived, or decellularized matrix) and most frequently include hydrogels [10,11]. Type I Collagen, hydrogels in 3D cell culture [12]. Patient-derived organoids (PDOs) have emerged as promising platforms for cancer research, with recent studies demonstrating their physiological and biological relevance for personalized medicine in HNSCC [13,14]. However, current 3D organoid protocols face diverse challenges, including variable success rates for their establishment, slow growth, small volumes and cell numbers, and the characteristic loss of stromal components during culture [15].

Previously, we described a simplified 3D cell culture method based on seeding both tumor- and stromal cells on top of ready-made Matrigel/type I collagen gels, instead of embedding the cells inside the gel. Embedding the cells homogeneously inside the gel, to which we refer as the “sandwich model”, sufficiently supports the growth of organoids from established cancer cell lines and also promotes tumor/stroma co-culture with CAFs [16]. However, the differential impact of these two approaches and their pros and cons on the composition of primary cultures of patient-derived tumor cell suspensions remains poorly investigated. In this study, different variations of seeding tumor cell suspensions were used and compared for patient-derived tumor cell cultures, as a) seeding the cells on top of the gel either more widely dispersed versus centered on a single spot, and b) by embedding the cells inside the gel, either dispersed or centered on a single spot. This is meant to impact cell densities, dispersion, and differential access to oxygen.

The main goals were as follows: a) to provide a rapid adaptation of patient-derived cell suspensions to 3D cell culture conditions after enzymatic tissue digest, b) to propagate these cells in simple 2D cultures with different growth media and to determine which media keep the original tumor heterogeneity best, and c) to generate a simple, robust, reproducible, informative, and physiologically relevant 3D platform for *in vitro* drug sensitivity assays in personalized medicine.

## Materials and Methods

### Patient information and tumor sample preparation

Tumor specimens were obtained during routine surgeries performed by two independent surgical teams at the Oncology Center of the Lublin Region (Centrum Onkologii Ziemi Lubelskiej, COZL) and the University Clinical Hospital No. 4 in Lublin (Uniwersytecki Szpital Kliniczny Nr 4 w Lublinie), Poland. Preoperative diagnostic evaluation was performed through biopsy to identify the tumor type. Tumor biopsies were delivered to our laboratory on the same day as the surgery. Written informed consent was obtained according to the local Ethics Committee (KE-0254/96/2020 and KB-0024/134/09/24) following the guidelines for institutional and national guidelines for the use of human material. Patient information is given in Table 1.

**Table 1:** Clinicopathological characteristics of four patients with oral and base of tongue malignancies.

| N° Patients | Age | Sex | Site | Histology | Size (cm) | LN Status | pTNM | Stage | Grade | Pre-op Tx | Surgical-Pathologic Findings | Recurrence |
| --- | --- | --- | --- | --- | --- | --- | --- | --- | --- | --- | --- | --- |
| 1 | 72 | Female | Right oral tongue | Keratinizing squamous cell carcinoma | 3.5 x 4.0 x 1.5 cm | 2/2 level III (+), 16 nodes (-), others reactive | pT3N2M0 | IVa | G1/G2 | None | Angioinvasion present; 2/2 LN level III positive; 0.1 cm deep margin | No local recurrence or distant metastasis |
| 2 | 41 | Male | Right mandible and floor of mouth | Keratinizing squamous cell carcinoma | 5.5 x 3.5 x 5.0 cm | 7/19 total positive | pT4aN2aM1 | IVc | G2-G3 | None | Perineural and angioinvasion; narrow margin; 7 LN positive (1/1, 4/4, 2/14) | Extensive lung metastases, local recurrence |
| 3 | 63 | Female | Base of tongue | Adenoid cystic carcinoma | 2.8 x 2.0 x 1.5 cm | 0/7 all negative | pT2N0M0 | II | Intermediate grade (cribriform + tubular), minor solid areas | None | Infiltrative growth into tongue, pharynx, tonsil; nerve and vessel resection; perineural invasion, Ki67 ~17% | No current evidence of distant metastasis |
| 4 | 62 | Male | Left oral tongue and floor of mouth | Keratinizing squamous cell carcinoma (recurrent) | 6.5 x 4.0 x 2.0 cm | 0/5 all negative | pT4aN0M0 | IVa | G2 | None | deep ulcerated mass; nodal necrosis; clear margins; R1 margin (deep) | Recurrent tumor, but no distant metastasis described |

### Preparation of patient-derived suspension cells and isolation of CAFs

Tumor samples were washed with DPBS (Thermo Fisher Scientific, USA), minced with a scalpel into pieces <1 mm, and dissociated using the gentleMACS™ Tissue Dissociator (Miltenyi Biotec, Germany) for more efficient homogenization before enzymatic dissociation. The homogenized samples were digested in a first round with 0.25%Trypsin-EDTA (Sigma-Aldrich, USA) for 30 minutes at 37 °C. The suspension was then filtered through a 100 µm cell strainer (Corning Falcon, USA). The filtered fraction, which is enriched in single cells and small clusters, was centrifuged at 900 RPM for 5 minutes and cryopreserved in 90% heat-inactivated FBS and 10%DMSO (both Sigma-Aldrich) in liquid nitrogen for later culturing. The undigested tissue remaining on the 100 µm filter was subjected to a second round of digestion with 0.25% Trypsin-EDTA for 60 minutes at 37 °C. This material was then sequentially filtered through 100 µm, 70 µm, and 40 µm strainers and cells centrifuged at 1500 RPM for 10 minutes. The resulting single-cell suspension was cultured in T25 flasks with Advanced DMEM/F-12 (Thermo Fisher Scientific), supplemented with 1% penicillin-streptomycin (10,000 U/mL), Primocin (100 μg/mL, Invivogen, USA), and 18% FBS under 5% CO₂ at 37 °C. This high-FBS condition was used to suppress the growth of squamous carcinoma cells and, instead, to promote CAF outgrowth. After 2 days, nonadherent cells were removed, and fresh medium was added. CAFs were expanded until 80–90% confluency, typically by passage 3, and subsequently frozen until used.

### IF staining and Western blot analysis for identification of CAF subtypes and preparation of CAF-conditioned medium

CAFs exhibiting distinct morphologies, as described in [5], were selected for IF staining using an anti-alpha smooth muscle actin (α-SMA) antibody (Abcam ab124964) and nuclear counterstaining with Hoechst 33342. The IF procedure was performed as previously described in [17]. Western blot analysis for CAFs was performed as previously described [18] with slight differences. Instead of 20 μg of total protein extract, 40 μg of protein extract was electrophoresed on 10% (w/v) and 5% (w/v) sodium dodecyl sulfate-polyacrylamide gel electrophoresis (SDS-PAGE) gels according to the size of the protein. The primary antibodies used were α-SMA (ab124964, Abcam, UK), anti-BMP4 (ab39973, Abcam), anti-Vimentin (sc-6260, Santa Cruz, USA), and anti-NOTCH3 (2889S, Cell Signaling Technology, USA). The secondary antibodies used were Horseradish peroxidase-conjugated (HRP) anti-rabbit IgG and anti-mouse IgG (both Cell Signaling Technology, #7074P2 and #7076P2). Pierce™ ECL Western Blotting Substrate (Thermo Fisher Scientific) was used to detect and quantify proteins and visualized by G: BOX Mini 9 Multi-Application Gel Imaging System (Labgene Scientific, Switzerland). After the identification and characterization, myCAFs at passage 4 were cultured in T-75 cell culture flasks with DMEM/F-12 containing 10%FBS until they reached 80-90%confluence. CAF-conditioned media were collected, centrifuged at 900 RPM for 5 min at 4 °C to remove non-attached cells and cell debris, and filtered with a Millex® MCE syringe filter 0.22 μm (Merck Millipore). The medium from C2-type CAFs and C1-type CAFs was mixed in a 1:1 ratio and was kept at −80°C until used in our tissue cultures.

### Preparation of hydrogels for direct 3D culture of patient-derived tumor cells

Matrigel (phenol-red free, Matrigel® Corning®; Germany) was used as a scaffold for 3D cultures in various combinations and ratios with type I collagen (rat tail type I collagen; Corning®) and Hyaluronic acid (HA), and buffered with HEPES (1M Gibco™ HEPES; Thermo Fisher Scientific). Hyaluronic acid was prepared as described previously with variations [19]: Briefly, 50 mg of Hyaluronic acid sodium salt (from Streptococcus, Thermo Fisher Scientific) was stirred with 5 ml of PBS containing Ca^2+^ and Mg^2+^ ions overnight and degassed. The pH was measured and adjusted to neutral pH 7.0. Gels were finally prepared at a final concentration of 2mg/mL Matrigel, 1mg/mL type I collagen, and 2%HA (w/v). A total volume of 600 µL gel mix was used for each well of a 6-well plate (Corning ® Costar ® TC-Treated Multiple) and allowed to polymerize for 2 hours in an atmosphere of 5% CO_2_ at 37 °C. Tumor cell suspensions were unfrozen and seeded on top of the prepared gels, at a density of 500.000 cells/well in our “media mix” composed of 1/3^rd^ of DMEM/F12 with 10% FBS, 1/3^rd^ of CAF-conditioned medium, and 1/3^rd^ of growth-factor-enriched Endothelial Cell Growth Medium 2 (ECM-2, PromoCell, Germany). After 2 days, non-attached, dead, and dying cells, cell debris, or undigested tissue fragments remaining were removed, the gel was washed with PBS, new medium was added, and the cells were cultured for up to 14 days. The morphological changes in primary cultures were visualized using the EVOS Cell Imaging Systems on days 2, 7, and 14.

### Transfer of primary tumor cultures from 3D to 2D conditions by gel digestion

To facilitate and simplify patient-derived tumor cell culture, we aimed to establish a straightforward and standardized 2D expansion protocol, followed by monitoring phenotypic changes that spontaneously emerge under different cell culture media. After 14 days in 3D culture, the Matrigel/Collagen/Hyaluronic Acid (MCH) gel was digested by gentle stirring with a solution of 3 mg/mL of Dispase II (Gibco™) at 37°C for 1 hour and subsequently centrifuged at 4 °C at 2000 RPM for 10 minutes. After the supernatant was removed, the pellet was washed with cold PBS and centrifuged at 4 °C at 1500 RPM for 5 minutes to remove any remaining gel fragments. Next, 100.000 cells/well were cultured directly on uncoated plastic surfaces using 5 different media: 1) high glucose DMEM supplemented with 2.5% FBS, 2) DMEM/F12 with 18% FBS; 3) “Media Mix 1” (ECM-2, CAF-conditioned medium, DMEM/F12 with 10% FBS as described above at a ratio of 1:1:1), 4) undiluted ECM-2 endothelial cell media with 2% FBS, and 5) “Media Mix 2” (50% ECM-2, 50% CAF-conditioned medium at 1:1 ratio). We generally found that growth-factor-supplemented ECM-2 endothelial media (with FGF5, EGF, IGF1, and VEGF) strongly supported the initial growth of primary tumor cell suspensions in both 2D and 3D, but that diluted ECM-2, mixed with other media, resulted in more differentiated cultures. Cells were passaged 3 times with the same medium. The entire process of cell culture adaptation from 3D to 2D is represented in Fig. 1.

**Figure 1:**
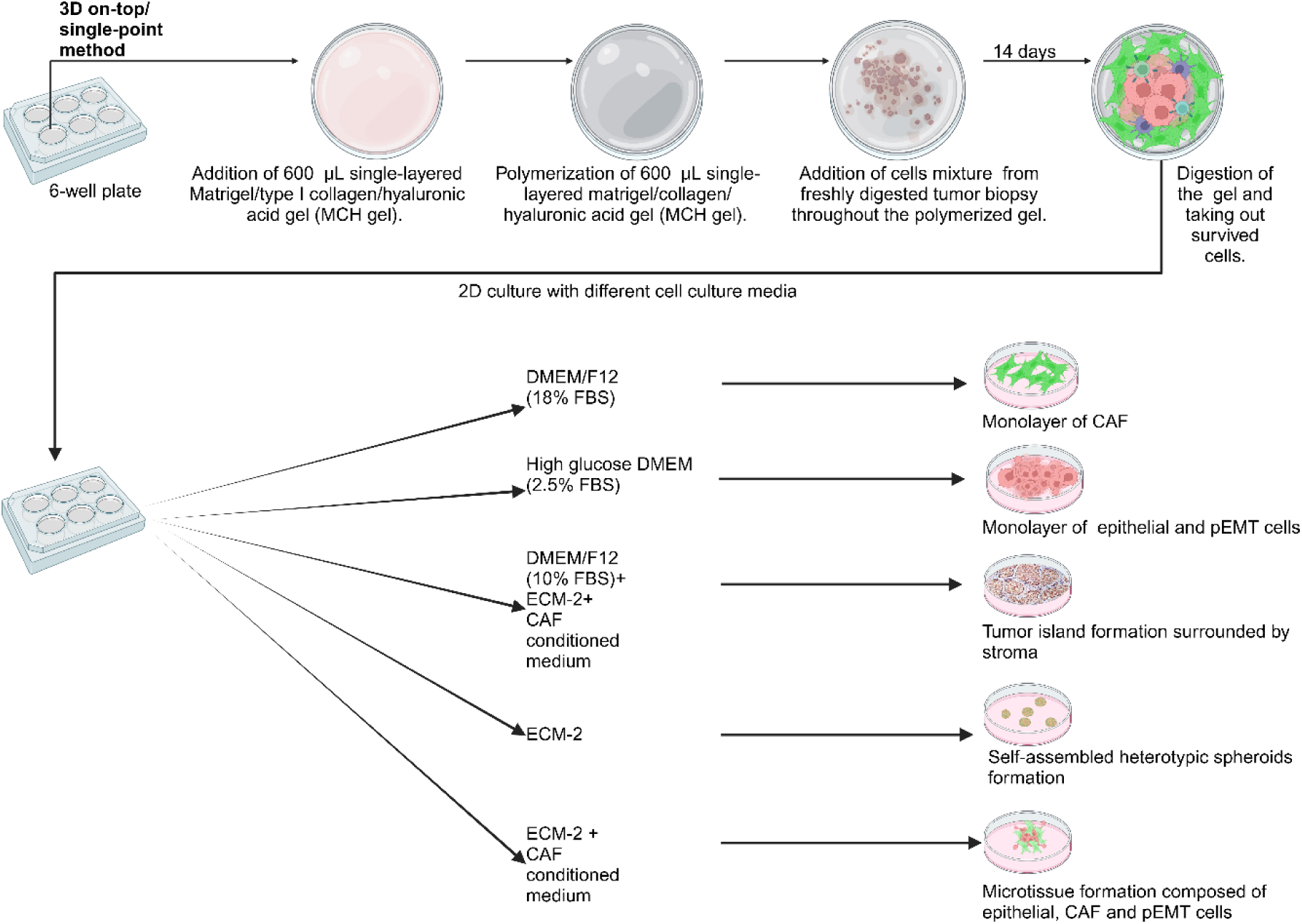
Primary culture of patient-derived tumor cells suspension after proteolytic dissociation of the tumor biopsy. Cryopreserved tumor cell suspension was cultured first on top of the MCH gel. After 14 days, the cells were extracted from the gel and propagated in 2D with different cell culture media. Different cell culture media resulted in the survival of various cell types and the spontaneous formation of tissue-like tumor structures (such as “tumor islands” surrounded by stromal areas).

After passage 3 in this “interim” type of 2D conditions, primary cell cultures with the 5 different media described above were divided into 3 groups, each cultured in 6-well plates. While the first group was passaged for further experiments, the second group, which contained 300.000 cells/well, continued to grow. When these cultures reached 80-90% confluency, the cultures were fixed with 4% PFA for subsequent IF staining. A third group of 2D cultures with 500.000 cells/well was directly used for extraction of RNA and quantitative real-time PCR (qRT-PCR).

### Immune Fluorescence (IF) staining for 2D cultures

IF staining was performed as previously described [17]. After final washing with PBS-BSA 3 times for 5 min, cells were imaged using a Nikon A1R-Si HD Confocal Microscope (Nikon Instruments Inc., USA). The 3D organoids obtained in 2D culture with 100% ECM-2 endothelial-cell media were visualized using a protocol for optical clearing, recently described for 3D structures [20]. The primary antibodies used were anti-E-Cadherin (ab76055, Abcam), anti-Vimentin (ab45939, Abcam), Anti-pan Cytokeratin (ab7753, Abcam), anti-beta-catenin (MA1-301, Invitrogen), and anti-alpha smooth muscle actin (α-SMA) antibody (ab124964). Goat anti-Mouse IgG (H+L) Cross-Adsorbed Secondary Antibody, conjugated with Alexa Fluor™ 647 (A-21235), and Goat anti-Rabbit IgG (H+L) Cross-Adsorbed Secondary Antibody, conjugated with Alexa Fluor™ 555 (A-21428, Invitrogen) were used as secondary antibodies. Hoechst 33342 dye was used for nuclear counterstaining.

### Quantitative real-time PCR (qRT-PCR) to monitor changes in mRNA gene expression in 2D cultures treated with different media compositions and serum concentrations

Total RNA was isolated using the RNeasy Mini Kit (QIAGEN, Netherlands). High-Capacity cDNA Reverse Transcription Kit (Applied Biosystems™, USA) was used for reverse transcription of 1 µg of total RNA. The SYBR® Premix Ex TaqTM reagent (TaKaRa, Dalian, China) was employed for qPCR analysis on an ABI PRISM 7500 real-time PCR system (Applied Biosystems, USA), using SYBR™ Green Universal Master Mix (Applied Biosystems) for qPCR analysis. Relative changes in mRNA expression were calculated using the 2−ΔΔCt method, normalized to GAPDH expression. Specific primers were used as follows:

GAPDH Forward: 5′-GAACGGATTTGGCCGTATTG-3′, Reverse: 5′-TTTGGCTCCACCCTTCAAG-3′;

CTNNB1 Forward: 5′-ATCCAAAGAGTAGCTGCAGG-3′, Reverse: 5′-TCATCCTGGCGATATCCAAG-3′;

PDGFR-β Forward: 5′-AGGACAACCGTACCTTGGGTGACT-3′, Reverse: 5′-CAGTTCTGACACGTACCGGGTCTC-3′;

VIM Forward: 5′-AGGAAATGGCTCGTCACCTTCGTGAATA-3′; Reverse: 5′-GGAGTGTCGGTTGTTAAGAACTAGAGCT-3′;

SNAI2 Forward: 5-ATCTGCGGCAAGGCGTTTTCCA-3′, Reverse: 5′-GAGCCCTCAGATTTGACCTGTC-3′;

SOX2 Forward: 5′-TACAGCATGTCCTACTCGCAG-3′, Reverse: 5′-GAGGAAGAGGTAACCACAGGG-3′;

TNC Forward: 5′-GGTACAGTGGGACAGCAGGTG-3′, Reverse: 5′-AACTG GATTGAGTGTTCGTGG-3′.

### Scaffold-based 3D cultures in µ-Plate 96-well plate

At passage 4, 2D-expanded cells were transferred again into 3D conditions to investigate the spontaneous differentiation and formation of tissue-like structures. For this purpose, we were using µ-Plate 96-well (Ibidi GmbH, Germany). Tumoroids’ morphology was evaluated based on three independent variables: The first factor was 1) gel composition, the second factor was 2) media composition, and the third factor was 3) differential oxygen availability and nutrient diffusion.

1. To evaluate the effects of different gel compositions, the 3D sandwich/single-point method was chosen initially for patient 1. The settings were prepared by first placing 30µL of the bottom gel into each well, allowing the gel to polymerize for 2h, followed by the addition of 10 µL unpolymerized gel on top. Before polymerization, 5.000 cells were injected with 5 µL of media into the center of the upper gel, where they formed a dense cell cluster. Four gel compositions were tested in DMEM/F12 with 10% FBS: a) 4 mg/mL pure Matrigel, b) 4 mg/mL Matrigel + 0.375mg/mL collagen I, c) 2mg/mL Matrigel + 0.75mg/mL collagen I, and d) 2mg/mL Matrigel + 1mg/mL collagen I
2. Next, we aimed to investigate whether different culture media could allow us to recapitulate the invasive features observed with tumoroids in 3D cultures. Here, we aimed to prevent undesired matrix degradation and contraction, combined with the adhesion of hyperactive CAFs at the bottom of the plates. Since this was preferentially observed with collagen or Matrigel/collagen type I mixed gels, we decided to switch to pure Matrigel for this purpose. Accordingly, cells from patient 1 were cultured using the 3D sandwich/single-point cell seeding method in 4 mg/mL pure Matrigel, testing the following media conditions: a) DMEM High Glucose with 2.5%FBS, b) DMEM/F12 with 10%FBS, c) full Endothelial Cell Growth Medium 2 (ECM-2), and d) Media Mix 2 (a 1:1 combination of full ECM-2 and CAFs-conditioned DMEM/F12 with 10%FBS). The IF staining was done as described previously [16].
3. After selecting 4 mg/mL pure Matrigel with Media Mix 2 as optimal conditions for tissue-like structures to emerge, patients 2 and 3 were assessed (in addition to patient 1 cells), using the same 3D cell culture conditions. Additionally, we aimed to address the impact of different cell seeding methods, which affect oxygen and nutrient diffusion, on tumoroid formation. Cell suspensions, transferred from 2D interim cultures of patients 1–3, were analyzed for this purpose (summarized and described in Fig. 2). The Morphology of tumoroids was evaluated using the complexity parameter, calculated as: Complexity = perimeter² / (4π × area) (adapted from [21]). ImageJ software was used for perimeter and area measurements [22], with a pixel-to-micrometer ratio of 0.80 pixels/μm.

**Figure 2:**
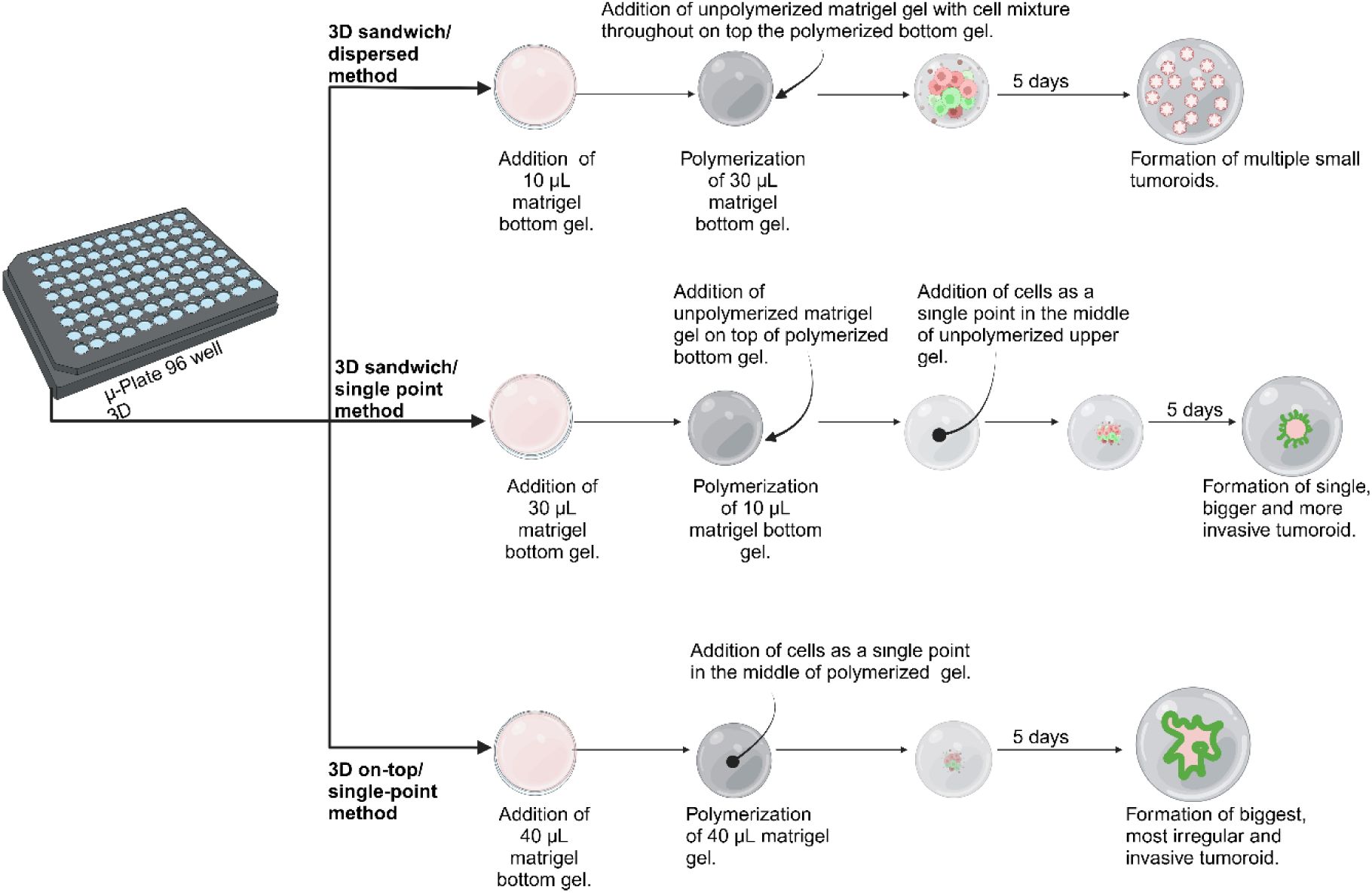
The cell seeding method affects the morphology and formation of tissue-like structures of tumoroids in 3D cultures. Three strategies were evaluated using equal cell numbers (5,000 cells/well): a) 3D sandwich/dispersed method: Cells were homogeneously embedded within a thick, unpolymerized upper gel layer, generating 3D cultures with reduced access to oxygen and nutrient diffusion. This growth condition resulted in small, slow-growing, uniformly distributed organoids with typically regular morphology. b) 3D sandwich/single-point method: Cell suspensions were injected into the center of a thin unpolymerized upper gel layer (but not on top), allowing improved semi-embedded formation of cells at high density, and improved access to oxygen and nutrients. This method led to the formation of a single, dense tumoroid. c) 3D on-top/single-point method: The cell suspension was placed as a droplet on top of the polymerized gel surface with minimal medium, resulting in high cell density, and offering maximal oxygen and nutrient exposure. This last approach produced the largest, most irregular, and rapidly growing tumoroids.

### Drug testing in 3D – phenotypic assays combined with WST-8 metabolic assay

For *in vitro* drug sensitivity testing in 3D tissue-like conditions (or as “tumoroids”), 5000 cells per well from patients 1-3 were grown in Media Mix 2 (full ECM-2 and CAF-conditioned medium mixed at a 1:1 ratio). Cell suspensions were trypsinized and cultured in single wells of the 96-well µ-angiogenesis plates (Ibidi GmbH, Munich) by directly seeding the cells on top of the 4 mg/mL Matrigel layer (protocol c) shown in Figure 2), and cultured for 5 days. Under these conditions, cell cultures rapidly formed large, complex three-dimensional multicellular aggregates, which rapidly developed strong aggressive/invasive properties, indicating high levels of cell motility. After 5 days, these tissue-like 3D aggregates were treated with 5 µM of Cisplatin, 5 µM of Notch pathway activator RIN-1, and 5 µM of Notch pathway FLI-06 for 2 days. The morphological changes of the complex were monitored and documented by phase-contrast microscopy using the EVOS Cell Imaging System. Changes in aggregate size (total area change by the drug were morphometrically measured and quantified by using Image J software. Cell Counting Kit 8 (WST-8/CCK8, ab228554, Abcam) was used to determine the number of living cells by Multimode microplate reader Infinite M200 PRO (Tecan, Switzerland).

### Statistical analysis

All statistical analyses were conducted with GraphPad Prism (version 1.4.1, GraphPad Software, USA). For monitoring changes in mRNA gene expression as a result of 2D and 3D cultures in different media conditions, two-way ANOVA was used. To evaluate the changes in complexity for the different cell seeding methods, repeated measures by one-way ANOVA were used. For growth area calculations and WST-8 results, two-way ANOVA with multiple comparisons mode based on a post hoc test was used. All data are presented in the form of mean ± SD and are the result of no less than 3 independently performed experiments. Statistical significance was determined as values of *P < 0.05, **P < 0.01, ***P < 0.001, ****P < 0.0001, and ns: not statistically significant.

## Results

### Identification of C1 and high α-SMA C2 subtypes of CAFs by IF staining and western blot

Two CAF subtypes were distinguished according to their morphology and proliferation rate [5]. C2-type CAFs, with rapid growth and less elongated shape, exhibited strong α-SMA staining. In contrast, elongated and slower-growing C1-type CAFs showed weaker α-SMA signals and lower cell density. Western blotting confirmed subtype-specific protein expression: C2-type CAFs expressed high α-SMA and NOTCH3 but low BMP4, while C1-type CAFs showed the opposite profile. Vimentin expression did not differentiate between the types. Representative IF and Western blot images are shown in Fig. 3.

**Figure 3:**
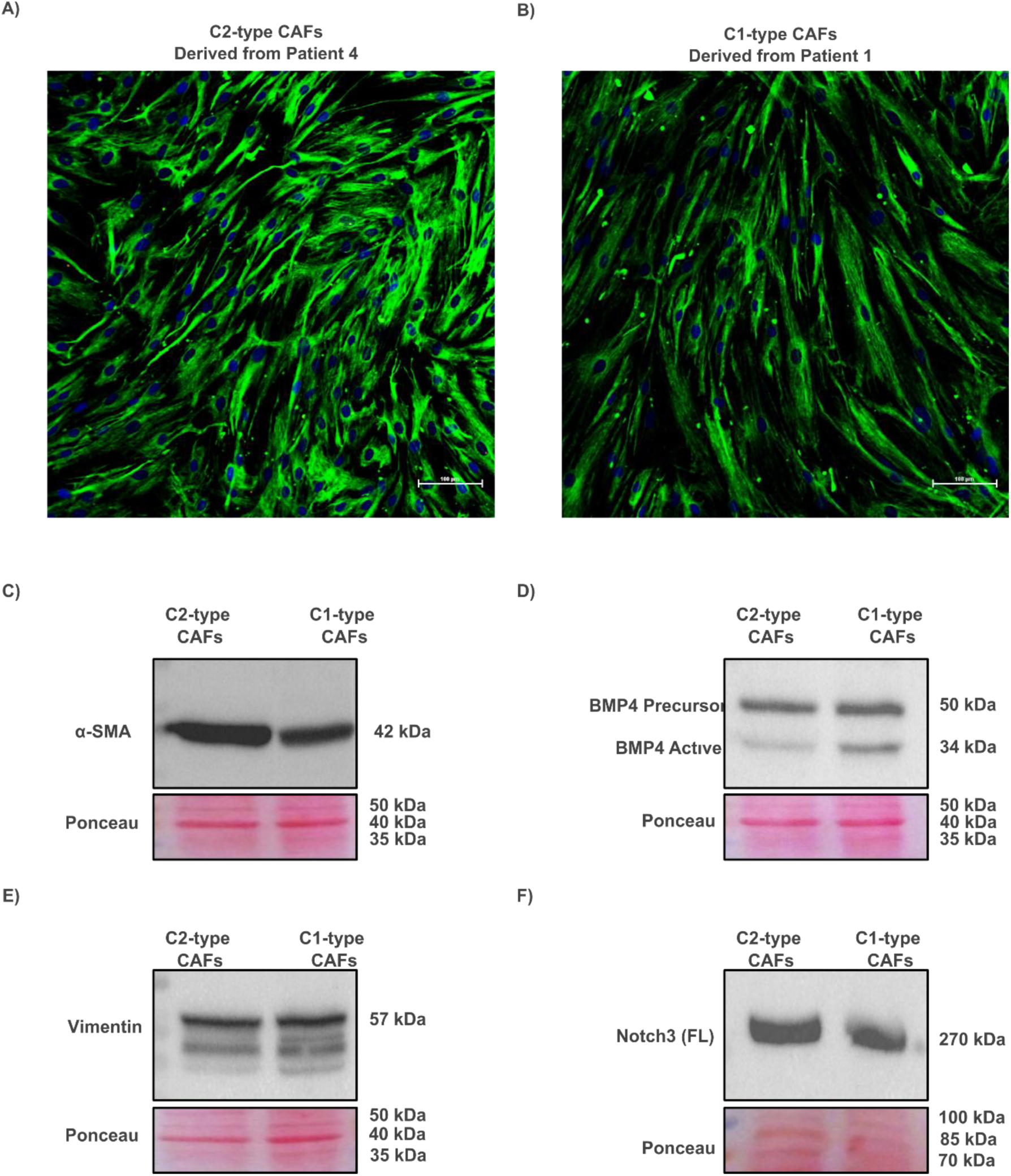
Characterization of CAF subtypes based on morphology and protein expression. A) IF staining of C2-type CAFs showing strong α-SMA expression. B) IF staining of C1-type CAFs with a weaker α-SMA signal. Nuclei were counterstained with Hoechst 33342. C–F) Western blot analysis of CAFs from two patients: (c) α-SMA, (d) BMP4, (e) vimentin, and (f) full-length NOTCH3. Ponceau S staining was used as a loading and transfer control.

### Culturing of patient-derived tumor cell isolates on top of Matrigel/type I collagen/hyaluronic acid (MCH) gel

Fig. 4 illustrates the adaptation process used for cell suspensions isolated from patient biopsies on top of the MCH gel over 2, 7, and 14 days. By day 2, all three patients formed small tumor cell aggregates, which differed in density and organization. In patient 1, the cells formed dense, string-like arrangements, which were later followed by steady expansion across the gel surface over time. Patient 2 exhibited the earliest signs of invasive behavior, with irregular aggregates showing outward cellular migration as early as day 2. These structures expanded and dispersed over time, forming multiple aggregates with a migratory pattern throughout the gel. Patient 3, derived from adenoid cystic carcinoma, exhibited aggregate fusion and compaction over time without extensive invasion into the surrounding matrix; they remained more localized, forming large, tumor-like nodules rather than scattering throughout the gel.

**Figure 4:**
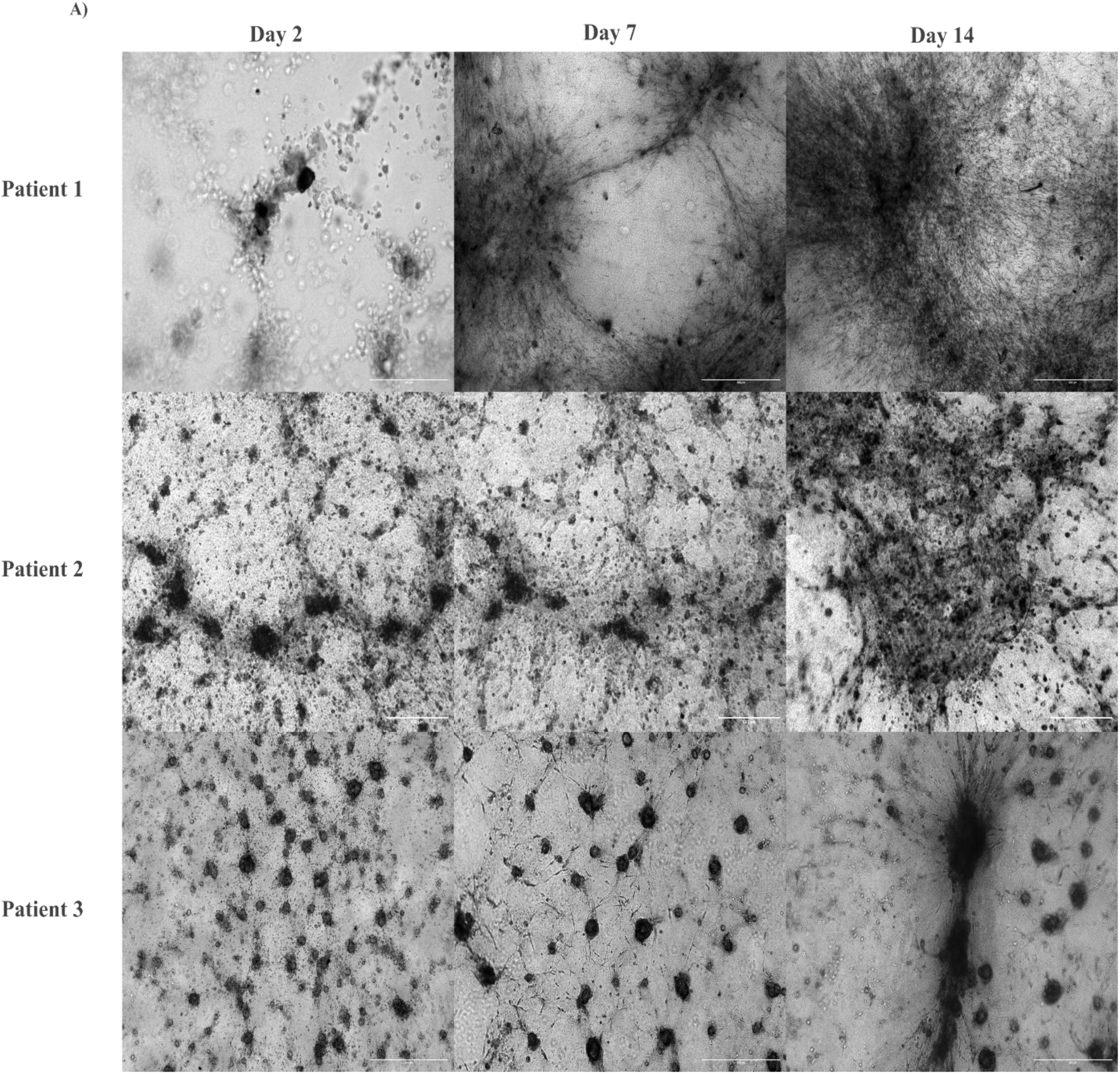
Time-dependent adaptation of tumor biopsy-derived cell suspensions on top of the MCH gel. Time-course images of tumor cell suspensions from three patients cultured on top of MCH gels at days 2, 7, and 14. Scale bar: 600 μm.

**Figure 5:**
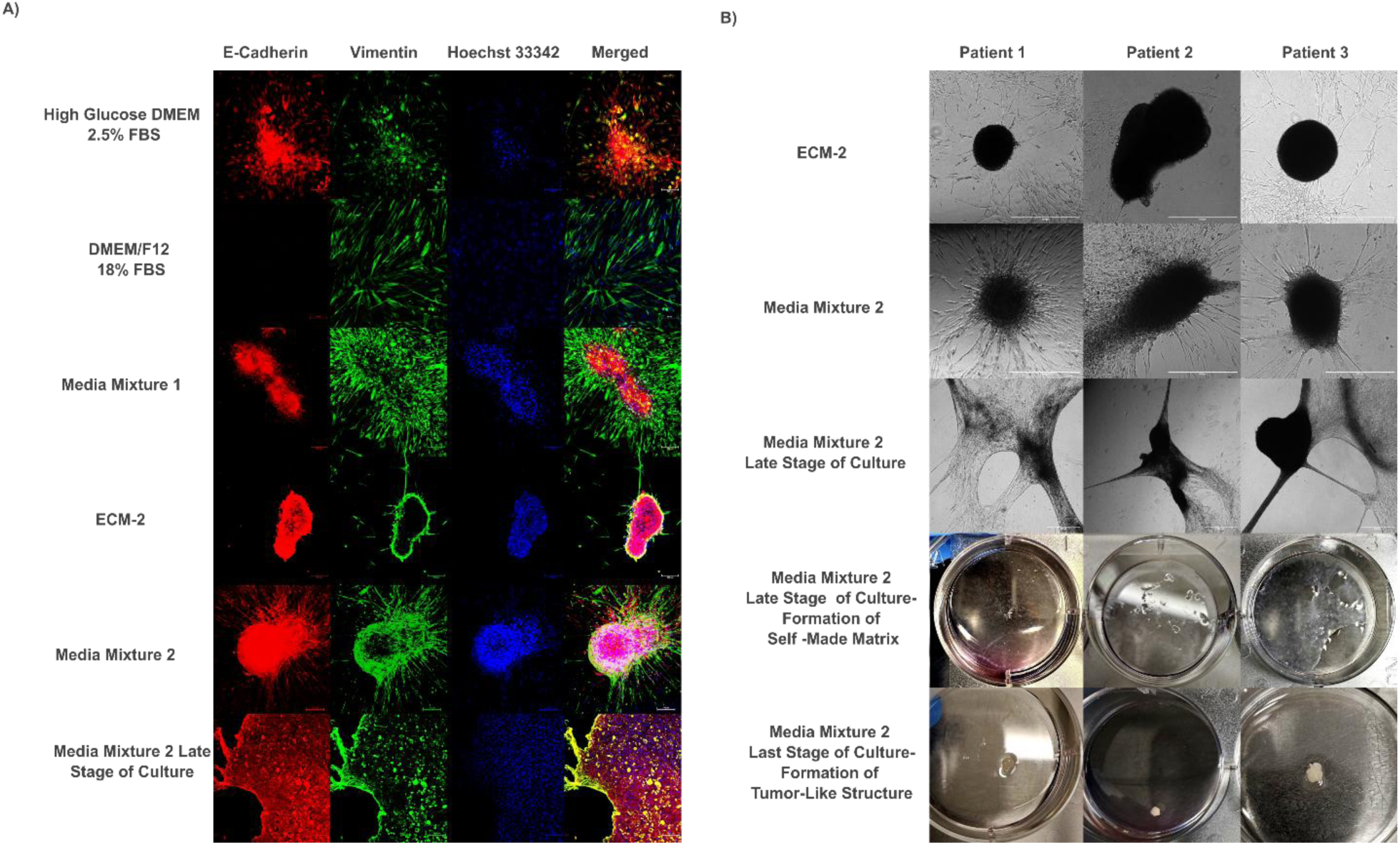
Culture media cause cell-type selection and 3D structure formation in 2D-expanded primary cultures. **A**) IF staining of patient 1 cultures grown on plastic under different media conditions, showing E-cadherin (epithelial), vimentin (mesenchymal), and nuclear counterstaining with Hoechst 33342. Simultaneous expression of E-cadherin and vimentin indicates partial epithelial-to-mesenchymal transition (pEMT). Scale bar is 100 μm. **B)** Brightfield and macroscopic views of cultures from patients 1–3 under spontaneous aggregation conditions. ECM-2 promoted the formation of nonadherent spheroid-like tumor aggregates, while Media Mix 2 supported adherent, invasive 3D structures. Scale bar is 600 μm.

**Figure 6:**
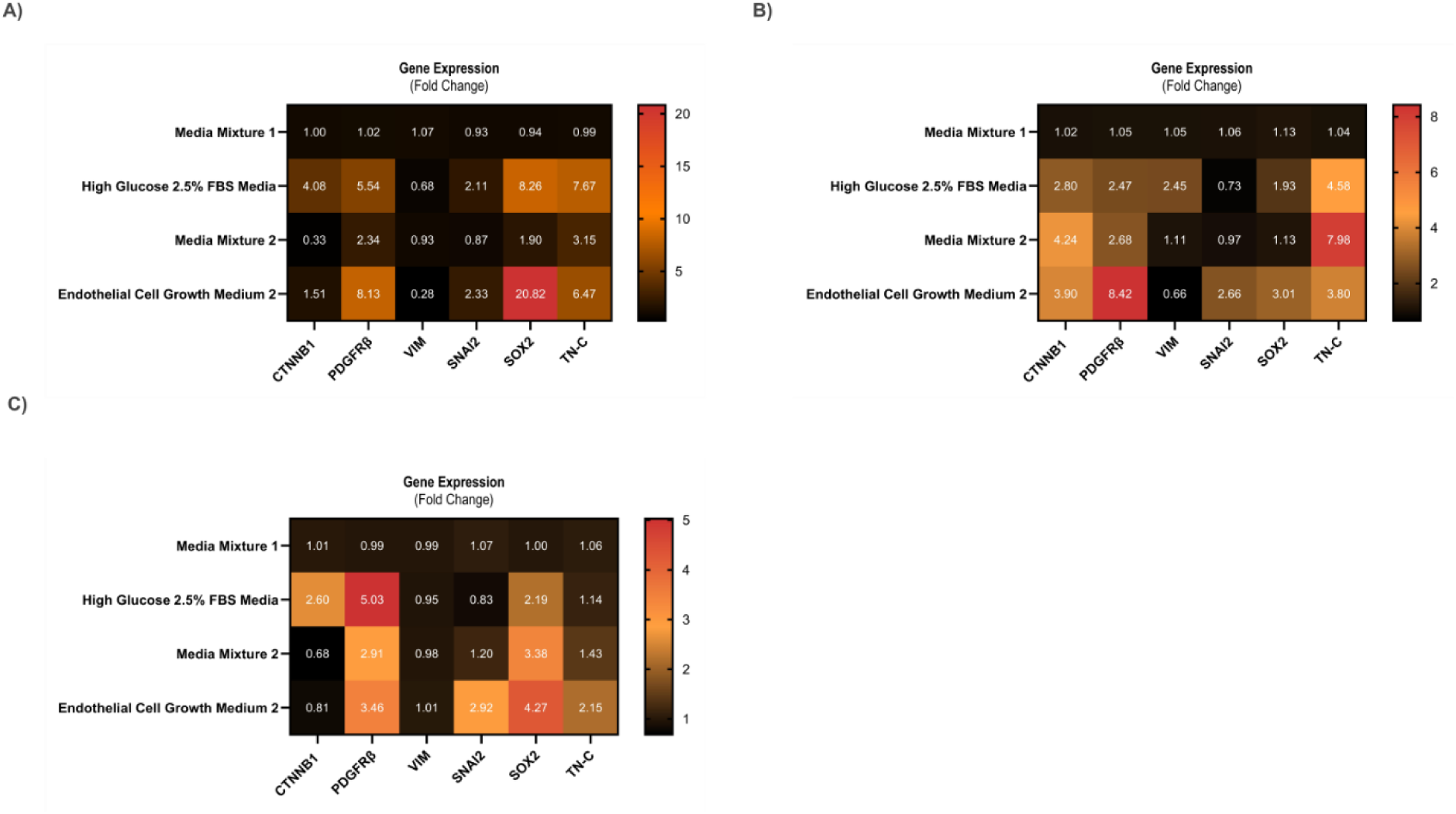
Different media have a profound effect on the expression of marker genes related to partial EMT and/or indicating stem cell characteristics. A) Relative gene expression for patient 1 cell cultures in 2D, exposed to different media as described in Fig. 5. B) Relative gene expression for patient 2 exposed to different media. C) Relative gene expression for patient 3 using the same media variations. Fold change values under 1 were considered as downregulated (black, *p ≤ 0.05) and ratios ranging from 1.93 to 20.82 as upregulated (red, n=3, *p ≤ 0.05), respectively.

### Variation of cell culture media leads to differential cellular composition in 2D tissue-like cultures

Media conditions strongly influenced the selection and retention of different cell types in 2D cultures, as demonstrated by strikingly different formation of multicellular structures and expression patterns of epithelial versus mesenchymal markers. Low-FBS DMEM media favored cell survival of tumor and pEMT cells, while high-FBS DMEM supported the expansion of CAFs. Media mix 1 (which retained 33% growth-factor-enriched ECM-2) effectively promoted the spontaneous formation of squamous carcinoma-like tissue histology with tumor islands surrounded by stromal zones. Full ECM-2 alone led to nonadherent spheroid-like aggregates composed of tumor and pEMT cells but did not support the retention of CAFs. In contrast, Media Mix 2 (which retained 50% full ECM-2 media and growth factors) fully supported the spontaneous aggregation of invasive spheroid-like aggregates that retained all cell types, including epithelial, pEMT, and CAF populations. These rapidly growing mixed 2D cultures eventually contracted and spontaneously formed thick, white/opaque, macroscopic tissue-like masses reminiscent of fibrosis, and completely detached from the plastic plates. Cells in these contracted and dense tissue-like masses retained viability (data not shown) but slowed down growth.

### qPCR analysis reveals different cell types and differentiation stages resulting from 2D tissue-like cultures treated with different media compositions

Primary tumor cell suspensions cultured in high glucose/low serum DMEM (2.5% FBS) expressed increased levels of beta-catenin (*CTNNB1), platelet-derived growth factor receptor beta (PDGFRβ)*, and SRY-box 2 (*SOX2)* mRNA across all samples. Vimentin (*VIM)* expression varied depending on the culture, showing both increases and decreases. Cell cultures grown in full ECM-2 media further upregulated *PDGFRβ*, *SOX2*, Snail Family Transcriptional Repressor 2 or Slug (*SNAI2)*, and tenascin C (*TN-C)*, while generally reducing VIM expression. Media Mix 2 induced moderate increases in *SNAI2, SOX2*, and *TN-C*, with relatively stable VIM expression. Gene expression pattern of epithelial vs mesenchymal markers in cell suspensions cultured in Media Mix 2 closely resembled those grown in Media Mix 1, with *CTNNB1* displaying variable regulation between cultures.

### Different growth conditions result in variable morphologies in scaffold-based 3D cultures

After selecting cells expanded in Media Mix as the standard source of tumor cells for further experiments, we started optimizing the 3D drug testing platform used for generating tissue-like tumoroids in 3D, investigating the impact of matrix, media composition, and topology of cell seeding.

To evaluate the impact of different hydrogel compositions on tumoroids’ morphology, cells were cultured in Matrigel with gradually increasing concentrations of collagen type I (Fig. 7A), using DMEM/F12 (10%FBS) as standard media, and the 3D sandwich/single-point method as the cell seeding method of choice This strategy offers relative moderate nutrient conditions (with low levels of growth factors) and moderate oxygen availability, due to slow diffusion into the gels. The use of pure Matrigel as a matrix results in the formation of small, non-invasive tumoroids with a small central area composed exclusively of squamous tumor cells, surrounded by a small number of fibroblasts with very low growth and motile activity. The incremental rise in type I collagen concentrations as ECM resulted in strikingly different morphologies of the tumoroids and functional activation of CAFs. Incremental addition of collagen eventually resulted in the rapid contraction of the entire 3D culture, followed by destruction of gel integrity after typically 5 days of culture. This was observed for all patients. The most stable 3D culture condition, which excluded gel contraction, was observed with pure Matrigel. However, we also observed that Matrigel did not support cell motility or invasiveness, and the resulting organoid-like structures were not representative of the original tumor histology and composition.
Next, we questioned if more invasive tumor/stroma co-cultures and multicellular, heterogeneous tissue-like tumoroids in Matrigel could also be obtained by switching between different media preparations (Fig. 7B), with more or less serum and growth factor supplements, or by adding fibroblast-conditioned media as a potent support for cell growth and spontaneous formation of tissue-like structures. Cell seeding was performed as described for the previous set of experiments using the 3D sandwich model and the single-point cell seeding method, which results in high local cell densities but also strongly supports cell survival and growth. High glucose DMEM media with low 2.5% FBS resulted in the formation of tumoroids that were largely or predominantly composed of tumor cells, but it does not support the growth of CAFs. In contrast, both ECM-2 media and Media Mix 2 (CAFs-conditioned media and ECM-2 1:1 mix), which are both enriched in growth factors such as EGF, IGF1, FGF2 and VEGF, resulted in the rapid formation of large and heterogeneous tumoroidss with strikingly invasive structures, with a large contribution of highly active CAFs in these sustained tumor/stroma cocultures. These structures were similar to those forming in hydrogels with high type I collagen composition, but did not require collagen for structure formation.
Lastly, we assessed the effects of oxygen and nutrient availability on each patient’s culture. For patients 2 and 3, the 3D sandwich/single-point method reproduced the most complex tumoroid morphology previously seen in patient 1. In contrast, the 3D sandwich/dispersed method resulted only in small, poorly developed, and slowly growing organoids, while the 3D on-top/single-point method produced the most complex, fast-growing, and irregular tumoroids, closely resembling native tumor architecture (Fig. 7C).

**Figure 7:**
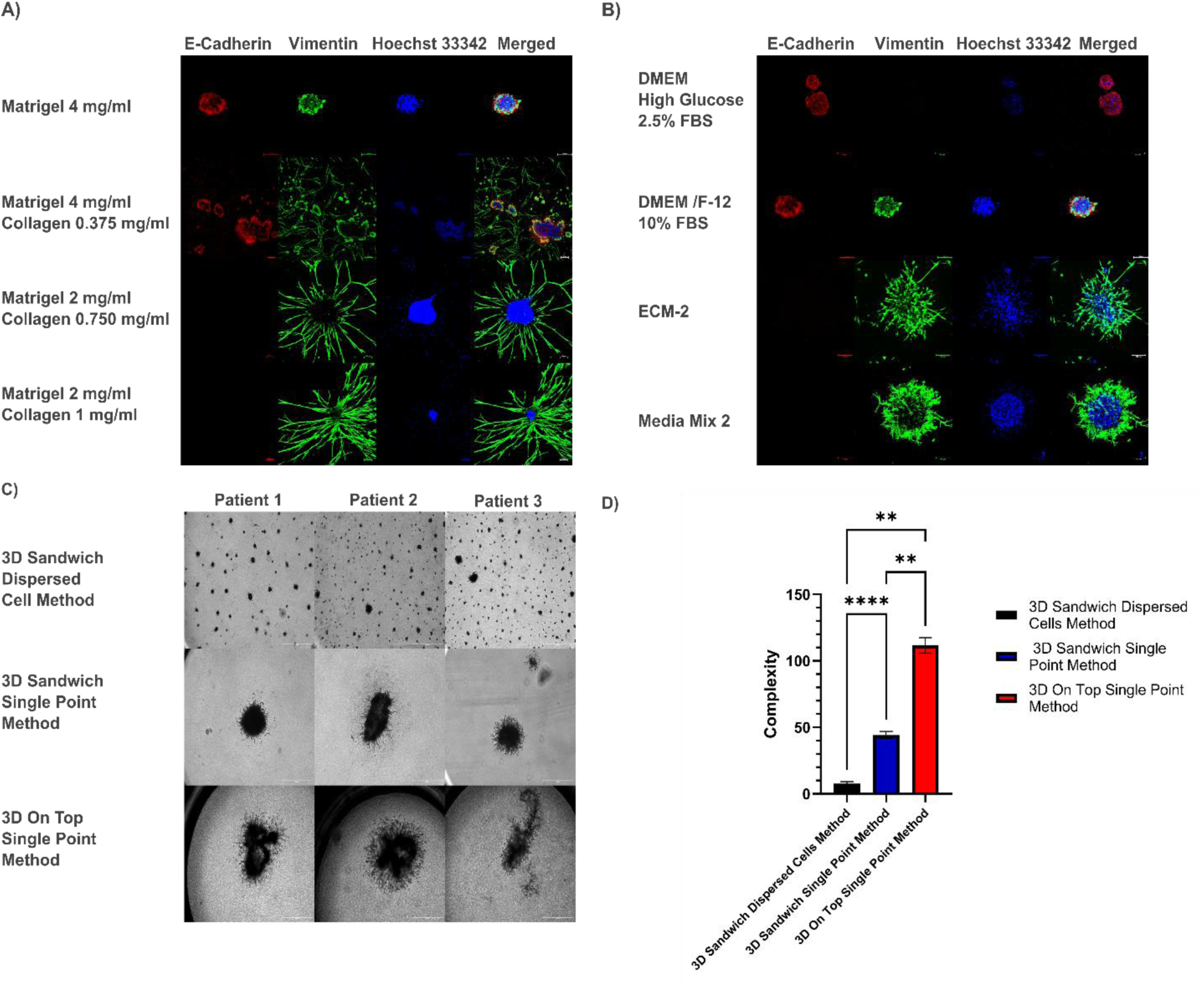
Gel composition, media, and seeding method have profound effects on the morphology of organoids forming in 3D cultures. A) Different gel composition affects the morphology of patient 1 cultures, which were treated with DMEM/F12 10%FBS and seeded with the 3D sandwich /single-point method. Scale is 100 μm. B) Phenotypic effects of tumoroids forming from patient 1 primary cell isolates on Matrigel as a scaffold, and different media as described above. Cells were seeded using the 3D sandwich/single point method on top of a single, uniform gel layer. Scale is 100 μm. C) The effects of different seeding methods on the morphology of patients 1, 2, and 3: All cultures contain Matrigel as a consistent hydrogel and use Media Mix 2 as media. The scale is 600μm. D) Complexity comparison for different cell-seeding methods. The complexity is shown as the mean ± SD (**p < 0.01, ****p < 0.0001; n = 3, one-way ANOVA).

### In vitro drug sensitivity testing on 3D cultures with patient-derived tumoroids

After selecting Matrigel as the hydrogel of choice, Media Mix 2 was chosen as the most reproducible and effective media composition, and the 3D on top single point method as the most effective seeding method. Using these conditions, tumoroids derived from patients 1, 2, and 3 were exposed to different small molecular weight inhibitors or drugs. Tumoroid morphology was monitored every day for 7 days (Fig. 8). We measured the total area of the tumoroids as an indicator for both tumor growth and invasion, and the readout of the metabolic WST-8 assay as a measure for cell viability. Notch pathway inhibitor FLI-06 resulted in the shrinking of the total area of tumoroids and concomitantly decreased cell viability for each patient. Concerning response to cisplatin, patients 1 and 3 were drug sensitive, and patient 2 was resistant. This differential cisplatin sensitivity likely reflects the clinical heterogeneity observed in patients: approximately 30-40% of the HNSCC patients show primary cisplatin resistance [1]. Notch pathway activator RIN-1 did not show any robust, reproducible effect on cell viability across patients. However, Notch pathway inhibitor RIN-1 notably restricted the invasive behavior of the cisplatin-resistant tumoroids derived from patient 2 (but not for patient 1 or 3).

**Figure 8:**
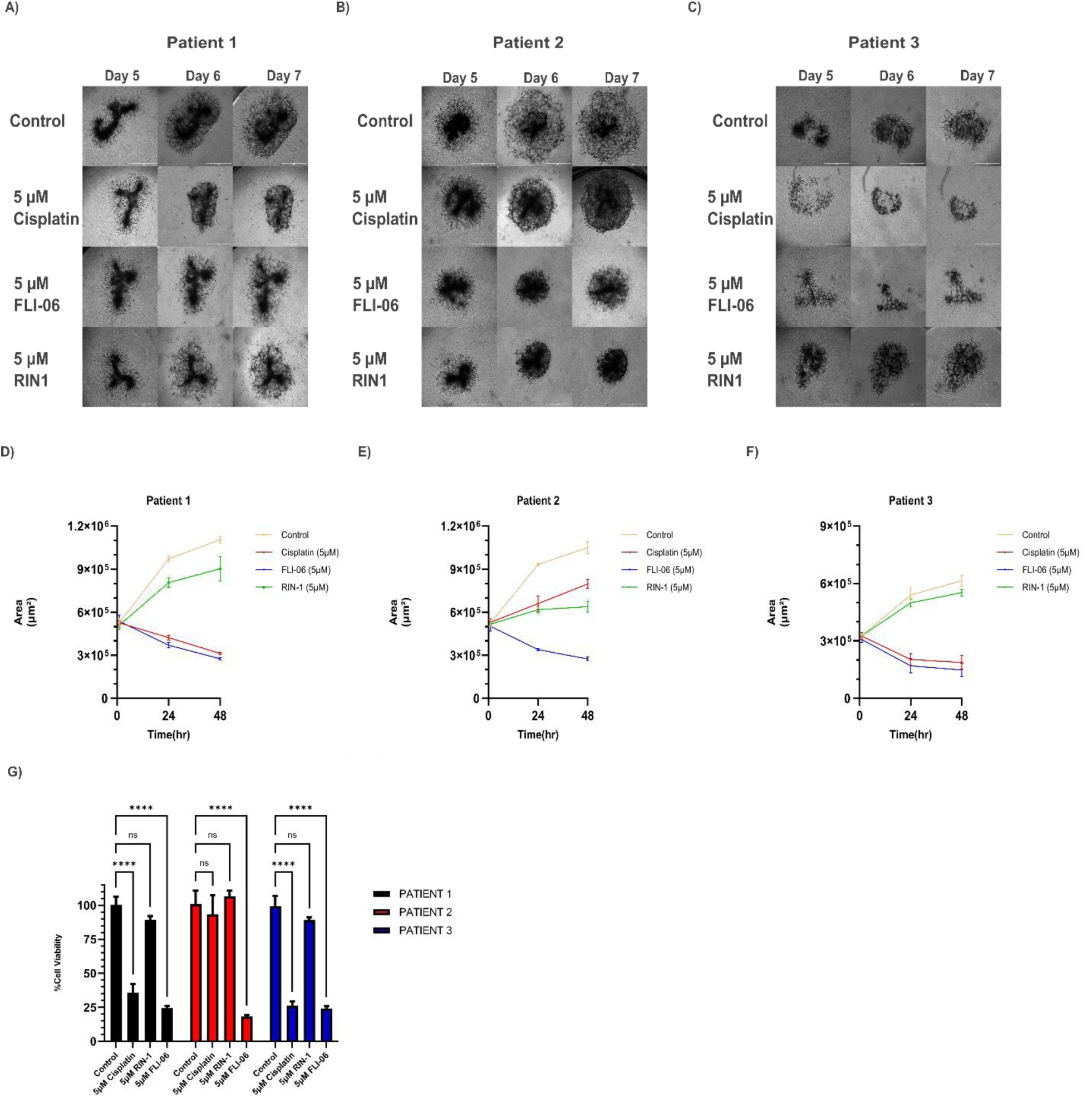
Different compounds differentially affect tumoroid growth and viability. A) After 5 days of seeding, tumoroids of patient 1 cells in media mixture 2 were exposed to 5 µM Cisplatin, 5 µM FLI-06, and 5 µM RIN1. The morphologic changes were observed on days 5, 6, and 7. B) and C) Same in vitro drug sensitivity test performed with cells from patients 2 and 3, respectively. D) Total area changes as observed after exposing patient 1’s cultures to different drugs. E) and F) Same in total area change for patients 2 and 3, respectively. Data represent the mean ± SD (*p < 0.05; n = 3, two-way ANOVA with multiple comparisons mode). G) WST-8 metabolic assay for patients 1, 2, and 3 cultures that were exposed to cisplatin, FLI-06, and RIN-1 at day 7. Data are mean ± SD (****P < 0.0001, ns: not statistically significant; n = 3, two-way ANOVA with multiple comparisons mode).

## Discussion

Recent systematic analyses reveal that among common cancer types, only 4.2% of head and neck cancer drug discovery studies utilize 3D in vitro models, highlighting a critical unmet need for physiologically more relevant HNSCC platforms [23]. Patient-derived xenograft (PDX) models are still considered the most realistic model system for in vivo drug sensitivity testing and personalized medicine. However, their prolonged establishment time, low efficiency, and replacement of human stroma with murine CAFs after 3–5 passages limit their accuracy in representing patient-specific tumor microenvironments [24]. A more widely used alternative to PDX is patient-derived organoid cultures (PDO), which provide a somewhat faster *in vitro* alternative, but PDOs are also often exceedingly slow and typically eliminate the non-tumorous microenvironment components [25]. In this study, we aimed to develop a rapid, animal-free *in vitro* platform starting directly from heterogeneous, complex patient-derived cell suspensions that can be sub-cultured without tedious experimental effort or requiring costly media compositions (as used for PDO). Furthermore, our approach readily allows switching from 3D to 2D cell culture methods, further decreasing technical complexity. Overall, our approach was designed to preserve tumor heterogeneity and to support in vitro chemosensitivity assays and personalized drug screening within a short time following patient operation. Our approach also addresses key specific limitations identified in current organoid/PDO technology. While traditional HNSCC organoids often lose stromal components during establishment, our scaffold-based system with CAF-conditioned media maintains tumor-associated fibroblasts throughout the entire period of 2D or 3D culture [13,15].

The critical challenge in processing fresh tumor tissue is maintaining cell viability while preserving the original tumor’s cellular heterogeneity after enzymatic digestion. One of the key factors is to select the right scaffold that thoroughly supports cell adhesion and survival. We developed a simple, defined hydrogel matrix combining Matrigel, type I collagen, and hyaluronic acid (HA), which together promote both viability and preserve the heterogeneity. Matrigel, derived from mouse Engelbreth-Holm Swarm (EHS) tumors, primarily consists of laminin (∼60%), collagen IV (∼30%), entactin (∼8%), and perlecan (∼2–3%), along with trace proteins and various growth factors [26]. Matrigel is well known to support epithelial tumor cell adhesion, growth, and survival, while minimizing apoptosis or anoikis after dissociation [25,27]. It has been shown to support epithelial tumor cells via integrin-mediated activation of PI3K/Akt signaling [28]. However, pure Matrigel is also known not to support mesenchymal cell types like CAFs. The addition of type I collagen, therefore, enhances the survival of not only the tumor cells isolated from HNSCC biopsies. Addition of HA promotes proliferation and invasiveness of both tumor and stromal cells via the RHAMM receptor, which is involved in cancer progression, chemoresistance, and metastasis [29].

The second key factor to promote growth and maintenance of *in vitro* cell cultures is the cell culture medium. While predefined organoid media support tumor cell survival, “organoid media” contains many additives and is specifically formulated to exclude stromal components [25]. Therefore, to specifically preserve much of the original cellular heterogeneity obtained from the dissociated biopsies, we formulated “Media Mix 1,” combining CAF-conditioned medium, ECM-2 endothelial growth medium, and DMEM/F12 medium supplemented with 10% FBS. Full ECM-2 contains low serum (2 %), but includes potent growth factors (EGF, FGF5, IGF1, and VEGF) that strongly support epithelial cell proliferation and invasion [30,31]. Last but not least, CAF-conditioned medium (based on DMEM/F12 with 5% serum) maintains both epithelial and mesenchymal phenotypes [32]. DMEM/F12 with 10%FBS was empirically found to support various cell types in parallel. The main drawback of fibroblast-conditioned media is that the composition is likely to be exceedingly complex but not defined. Furthermore, the addition of equally undefined FBS provides a natural cocktail of factors essential for cell attachment, growth, proliferation, and survival [33], but its composition is also unknown, undefined, and likely variable.

A third key factor for successful tissue-like tumoroid culture protocols was found to be the topology of cell seeding, in addition to cell density and volumes used. Differential seeding topology (“on top” versus embedded or partially embedded) directly affects oxygen and nutrient availability. Topical cell seeding likely provides conditions similar to the “air-liquid interface,” which is known to support cell differentiation and has been successfully used for many applications. Insufficient oxygen and nutrient diffusion immediately after tissue dissociation and embedding into hydrogel scaffolds can lead to increased cell death, apoptosis, and the release of debris, free DNA, metabolites, and active enzymes, along with pH shifts [34]. In contrast, improved access to oxygen and nutrients, combined with growth factor-rich media, supports the survival and rapid establishment of primary cell and tissue cultures from patient-derived isolates.

After an initial 3D adaptation on MCH gels, cells were further expanded on 2D plastic to accelerate the cell growth, provide a larger number of cells for subsequent experiments, while still largely preserving the original tumor heterogeneity. This includes the stromal CAFs, key regulators of tumor progression and epigenetic modulation [35,36]. Five different media conditions or mixes were tested for their ability to support epithelial tumor cells, CAFs, and partial EMT or pEMT cells, identified by IF staining for E-cadherin (CDH1), vimentin (VIM), and their co-expression [37,38]. High-FBS DMEM favored CAFs but induced terminal differentiation in tumor cells [16], while low-FBS high-glucose DMEM supported tumor and pEMT-like cells [39,40]. Media Mix 1, with intermediate FBS content and added but reduced levels of growth factors, preserved both compartments and promoted spontaneous differentiation of tumor/CAF co-cultures, resulting in the formation of tissue-like structures distinguished as tumor islands, formed of squamous carcinoma cells, surrounded by stromal structures, thus faithfully mimicking squamous carcinoma histology [41].

In contrast, pure 100% ECM-2 endothelial-cell medium with reduced 2% serum likely led to overstimulation of some of the cell types in question, but repressed others, and resulted in the formation of nonadherent spheroid-like aggregates that were rich in epithelial/pEMT cells but excluded CAFs. At a later time point, these cultures also tended to detach from the cell culture plates, contract, and form fibrotic, dense cell masses. In contrast, Media Mix 2, again with reduced concentrations of growth factors, successfully maintained epithelial, CAF, and pEMT populations. Simultaneously, Media Mix 2 also promoted the production and secretion of ECM components and resulted in the formation of adherent, invasive structures that matured into dense, translucent tissue-like masses.

These properties and the cellular composition emerging or retained in these diverse cell culture conditions were further investigated using qRT-PCR analysis. For this purpose, we were checking the expression of epithelial (CTNNB1), mesenchymal (VIM), stemness (SOX2, PDGFRβ), and pEMT-related (SNAI2, TN-C) markers [6,42,43] in the various cell cultures of tumoroids. High-glucose DMEM induced expression of epithelial markers CTNNB1, PDGFRβ, and SOX2 across all patients, likely due to low FBS stress [44]. Full ECM-2 endothelial medium further elevated PDGFRβ, SOX2, TN-C, and SNAI2 expression, but resulted in reduced VIM in two patients. This likely indicates reduced numbers of CAFs in these cultures, or marked CAF exclusion, while at the same time, ECM-2 promoted the formation of round, stem-cell-like spheroids. Although PDGFRβ expression may indicate mesenchymal cell properties, its increase observed here can also indicate strongly promoted tumor cell plasticity that often comes in combination with increased stemness characteristics [43]. Notably, only Media Mix 2 enabled all three major cell populations or cell lineages to co-exist and support each other, also leading to increased levels of tumor cell invasion that were different between the patients and also responded differently to NOTCH modulators and cisplatin. Expression of CTNNB1 was increased only in tumoroids from Patient 2 (tumor-driven), but decreased in the tumoroids from cell suspensions of Patients 1 and 3 (CAF-driven). Unlike ECM-2 media, Media Mix 2 did not induce excessive stemness, and unlike Media Mix 1, it facilitated matrix remodeling and invasive outgrowths. Thus, Media Mix 2 was selected as the most suitable condition for downstream 3D assays, preserving some of the original cell heterogeneity while avoiding unwanted lineage drift or loss of some cell types.

We further optimized in vitro 3D growth conditions for the use of chemosensitivity testing by developing a fast, reproducible experimental model system that preserves tumor heterogeneity and invasiveness, offering a more practical, cost-effective, and rapid alternative to PDX models. While the addition of type I collagen enhanced the tumor–CAF crosstalk, it also triggered gel contraction and the culture’s failure due to CAF hyperactivation, likely via α2β1/α11β1 integrins and downstream FAK signaling [45]. To maintain stability of the in vitro culture setup, we used Matrigel as the scaffold. Media Mix 2 proved most effective in supporting the formation of invasive, heterogeneous tumoroids. We also evaluated three distinct seeding strategies: dispersed embedding, single-point, and on-top single-point seeding. Of these, the on-top/single-point method generated the most complex, irregular tumoroids—likely due to improved oxygenation, nutrient diffusion, and cell–cell communication. These morphological differences were quantified using the complexity parameter (perimeter² / 4π × area), which reflects shape irregularity and heterogeneity [21,46]. In contrast, dispersed embedding led to uniform but poorly proliferating structures with limited complexity and failed CAF support, likely due to restricted diffusion of soluble factors and oxygen [47,48].

Finally, we used this optimized tissue-like tumoroid model system to test for chemosensitivity against clinical drugs (cisplatin) and next-generation Notch modulators (FLI-06 and RIN-1). Among the tested compounds, the Notch pathway inhibitor FLI-06 [9] was the most effective, reducing cell viability across all three patient-derived tumoroids. This is likely due to its ability to induce apoptosis in cells with active Notch signaling and to cause cell cycle arrest in the G0/G1 phase. This is consistent with previous findings on Notch inhibitors in other solid tumors [49]. In addition, our recent work by Czerwonka et al. demonstrated that FLI-06 induces G0/G1 arrest and apoptosis across multiple HNSCC models, supporting our findings [9]. Cisplatin showed variable efficacy: while two cultures were sensitive, patient 2’s tumoroids were resistant. In contrast, the Notch-activator RIN-1 [8] did not significantly reduce viability or limit expansion in either Patient 1 or 3, but selectively restricted further growth of Patient 2’s cisplatin-resistant tumoroids. These findings point to a possible association between Notch signaling activity and cisplatin resistance, but they warrant further investigation.

### Study Limitations and Future Directions

Our study’s primary limitation is the small patient cohort (n=4, with n=3 for drug testing), which limits statistical power for biomarker discovery. However, our findings provide important proof-of-concept for the platform’s feasibility and demonstrate patient-specific drug responses that warrant validation in larger cohorts. Additionally, we did not obtain treatment outcome data for clinical correlation analysis, which represents an important future direction. Recent clinical trials, such as the SOTO study, are investigating organoid-guided treatment selection in HNSCC patients and may provide frameworks for future validation studies [50].

## Conclusions

We developed a robust and reproducible scaffold-based 3D culture platform for personalized medicine in head and neck cancer (HNC). Starting from fresh patient tumor biopsies, our system enables rapid *in vitro* adaptation, expansion, and reconstruction of tumor microtissues with persistent heterogeneity, including stromal and pEMT components. Through the optimization of hydrogel composition, media, and oxygen exposure, we established a functional, patient-specific drug testing system. This clinically connected platform bridges surgical oncology and laboratory research, providing a practical and time-efficient tool for guiding individualized therapeutic decisions and personalized medicine.

## Data availability

The authors confirm that the data supporting the findings of this study are available within the article. Raw data that support the findings of this study are available from the corresponding author upon reasonable request.

## Abbreviations

HNC: Head and Neck Cancer
HNSCC: Head and Neck Squamous Cell Carcinoma
CAFs: Cancer-Associated Fibroblast(s)
myCAFs: Myofibroblastic Cancer-Associated Fibroblast(s)
pEMT: Partial Epithelial-to-Mesenchymal Transition
PDOs: Patient-derived organoids
TME: Tumor Microenvironment
ECM: Extracellular Matrix
ECM-2: Endothelial Cell Growth Medium 2
HA: Hyaluronic Acid
MCH: Matrigel/ Type I collagen/ Hyaluronic Acid
α-SMA: Alpha Smooth Muscle Actin
BMP4: Bone Morphogenetic Protein 4
TN-C: Tenascin-C
CTNNB1: Catenin Beta 1
SNAI2: Snail Family Transcriptional Repressor 2 (Slug)
SOX2: SRY-Box Transcription Factor 2
PDGFRβ: Platelet-Derived Growth Factor Receptor Beta
IF: Immunofluorescence

## Funding

This research was funded in whole or in part by the National Science Centre, Poland, grant numbers (UMO-2020/37/B/N24/03920) and (UMO-2021/41/B/NZ7/03786OPUS19).

## Author Information

### Authors and Affiliations

Department of Biochemistry and Molecular Biology, Medical University of Lublin, 20-093 Lublin, Poland

Alinda Anameriç, Emilia Reszczyńska, Andrzej Stepulak, Matthias Nees

Department of Plant Physiology and Biophysics, Institute of Biological Sciences, Faculty of Biology and Biotechnology, Maria Curie-Skłodowska University, Akademicka St. 19, 20-033 Lublin, Poland

Emilia Reszczyńska

Otolaryngology Department of Saint Jan of Dukla Oncology Centre of the Lublin Region, Jaczewskiego 7 Street, Lublin, Poland

Tomasz Stankiewicz

Department of Otolaryngology and Laryngeal Oncology, Medical University of Lublin, Jaczewskiego 8 Street, 20-954, Lublin, Poland

Adrian Andrzejczak

## Contributions

Conceptualization: A. Anameriç, M.Nees, E. Reszczyńska, Methodology: A. Anameriç, E. Reszczyńska, Investigation: A. Anameriç, Formal analysis: A. Anameriç, E. Reszczyńska, M. Nees, Data curation: A. Anameriç, Resources (Patient material and surgical procedures): T. Stankiewicz, A. Andrzejczak, Supervision: A. Stepulak, M. Nees, Writing – original draft: A. Anameriç, M. Nees, Writing – review & editing: A. Anameriç, E. Reszczyńska, M. Nees, Visualization: A. Anameriç, Funding acquisition: M.Nees, A. Andrzejczak

## Ethics declarations

This study was approved by the Bioethics Committee of the Medical University of Lublin, Poland, and conducted in accordance with the ethical standards of the institutional and national research committees, and with the 1964 Declaration of Helsinki and its later amendments. The project was carried out under the ethical approval titled “Terapia eksperymentalna nowotworów głowy i szyi w warunkach doświadczalnych z użyciem hodowli pierwotnych” (Approval No. KE-0254/96/2020), with an amendment approved by resolution KB-0024/134/09/2024 on September 19, 2024. Patient-derived tissue samples were collected under this ethical approval, and all procedures were performed in accordance with institutional and national ethical standards.

## Competing interests

The authors declare no competing interests.

## Notes

### Competing Interest Statement

The authors have declared no competing interest.

